# Identification of plant exclusive lipid transfer SMP proteins at membrane contact sites in Arabidopsis and Tomato

**DOI:** 10.1101/2022.12.14.520452

**Authors:** Carolina Huercano, Francisco Percio, Victoria Sanchez-Vera, Jorge Morello-López, Miguel A Botella, Noemi Ruiz-Lopez

**Author notes:** Corresponding author: Noemi Ruiz-Lopez. These authors contributed equally to this work (Carolina Huercano, Francisco Percio). **Email addresses:** Carolina Huercano Francisco Percio Vargas Jorge Morello López Miguel A Botella Victoria Sánchez Vera Noemi Ruiz-Lopez.

## Abstract

Membrane contact sites (MCS) are regions where two membranes of different organelles are close but not fused; they coordinate non-vesicular communication between organelles and are involved in a wide variety of physiological functions, including membrane lipid homeostasis. Amongst proteins localized at MCS are those containing a lipid transport domain known as synaptotagmin-like mitochondrial-lipid binding protein (SMP), being the mammalian Extended Synaptotagmins, the yeast Tricalbins and the plant Synaptotagmin 1 (SYT1) the best SMP proteins characterized so far. They are all localized at endoplasmic reticulum-plasma membrane contact sites (ER-PM CS). We have carried out *in-silico* genome-wide identification of genes encoding SMP proteins in Arabidopsis and tomato. We have identified the plant exclusive NTMC2T5 proteins as ER-chloroplast CS components which make them extremely interesting as the route for lipid trafficking into and out of chloroplasts remains unknown. Additionally, *NTMC2T5* over-expressions caused a significant clustering of chloroplast around nucleus. Moreover, SYT6, NTMC2T6 and TEX2 have been identified as ER-Trans-Golgi Network CS proteins. These proteins associated between them and with the exocytosis related proteins VAMP721 and VAMP727. Since the functional roles of many of these genes are unknown, this gene collection provides a useful resource for future studies.

**HIGHLIGHT:** Plant exclusive lipid transport proteins were identified at membrane contact sites. SYT6, TEX2 and NTMC2T6 proteins are localized at ER-TGN. NTMC2T5 proteins are localized at ER-Chloroplast and induced chloroplast-nucleus clustering.

## INTRODUCTION

A distinctive feature of eukaryotic cells is their compartmentalization into organelles that perform specific functions. In consequence, protein function is crucially defined by its subcellular localization, as organelles exhibit distinct chemical environments and interaction partners. In the 1950s, it was discovered these organelles were not fully independent, since there were regions where the membranes of the two organelles were very close (typically 10–30 nm) without fusing (Bernhard and Rouiller, 1956). Nowadays, these areas are named membrane contact sites (MCS) and they have been extensively studied in the last 10-15 years. Some recent reviews have summarized the set of characteristics owned by these MCS (Jain and Holthuis, 2017; Wu *et al*., 2018; Scorrano *et al*., 2019): (1) the presence of tethering forces that arise from protein–protein or protein–lipid interactions, (2) the lack of membrane fusion at these sites, (3) the presence of specific proteins/lipids that are enriched at these MCS and (4) the fact that MCS accomplish specific functions including the specific bidirectional transport of molecules, the transmission of signalling information and the positioning in *trans* of some enzymes. Although MCS share these common elements, there is considerable diversity in the shape, protein/lipid compositions, stability, or the subcellular localization across different MCS. Live-cell microscopy, electron tomography and the development of unique fluorophores have allowed the identification of MCS between the ER and mitochondria, Golgi, endosomes, peroxisomes, lipid droplets, or the PM among others (Scorrano *et al*., 2019). Importantly, these observations are being supported by the identification/characterization of proteins involved in membrane tethering and/or functions of these MCS.

To date, several families of proteins which have the ability to transfer lipids at MCS (between membranes of the two organelles) have been discovered. For example, the ERMES complex was identified at ER–mitochondria CS and includes Mmm1 (a protein anchored at the ER), Mdm12 (cytosolic) and Mdm34 (which associates with the outer mitochondrial membrane) (Kornmann *et al*., 2009), all of them containing a synaptotagmin-like-mitochondrial-lipid binding protein (SMP) domain. These proteins belong to the TULIP superfamily, as the SMP domain is a barrel of about 200 aminoacids long whose interior is lined by hydrophobic residues creating a hydrophobic channel which transfers glycerolipids (AhYoung *et al*., 2015) between the associated membranes. SMP domains are conserved across species, from yeast to humans and they are typically found in intracellular proteins that locate at MCS (Kornmann *et al*., 2009; Reinisch and de Camilli, 2016; Jeyasimman and Saheki, 2020). Several proteins have been identified in the SMP-like family. In yeast, in addition to the proteins in the ERMES complex mentioned above, Tricalbins (Tcb1, Tcb2, and Tcb3), and Nvj2 are also SMP proteins. Tricalbins are critical tethering factors at ER–PM CS (Toulmay and Prinz, 2012) and recent studies have shown that Tcb3 is able of transferring phospholipids between the two membranes in the presence and even in the absence or Ca^2+^ (Qian *et al*., 2021). Nvj2 is a SMP protein which during ER stress, increases at ER–Golgi CS. This protein facilitates ceramide transfer from the ER to the Golgi complex, preventing the accumulation of ceramides under these conditions (Liu *et al*., 2017).

In animal cells, a total of four different groups of SMP proteins have been identified: E-Syt, TMEM24, TEX2 and PDZ8. The three extended-synaptotagmins (E-Syt1, E-Syt2 and E-Syt3) bring together the ER with the PM. E-Syt have an N-terminal transmembrane (TM) region that anchor them to the ER, followed by the SMP domain and from three to five C2 domains in the C-terminal side for PM binding (Giordano *et al*., 2013; Idevall-Hagren and de Camilli, 2015). E-Syt and Tcb form homo- or heterodimers due to dimerization of their SMP domains. In *vitro* analyses of E-Syt have shown that E-Syt transfer glycerolipids (regardless of their head groups) between membrane bilayers (Saheki *et al*., 2016*a*; Yu *et al*., 2016; Bian *et al*., 2018). Additionally, TEX2 and Transmembrane protein 24 (TMEM24) are also mammalian SMP proteins. TMEM24 also localizes at ER-PM CS and TEX2 at tubular ER interacting will the Golgi Apparatus (Wang *et al*., 2017; Sun *et al*., 2019). And the most recent discovered mammalian SMP protein is PDZ8 which has been shown to act together with TEX2 to prevent the build-up of PI(4,5)P_2_ on endosomal membranes (Raczkowski *et al*., 2017; Gao *et al*., 2021; Jeyasimman *et al*., 2021).

In plants, there is a bigger number of SMP proteins. Five E-Syt orthologues (SYT1–5) have been identified in *Arabidopsis thaliana*, being SYT1 the best characterized so far (Schapire *et al*., 2008; Yamazaki *et al*., 2008, 2010; Lewis and Lazarowitz, 2010; Uchiyama *et al*., 2014; Levy *et al*., 2015; Perez-Sancho *et al*., 2015; Kim *et al*., 2016; Siao *et al*., 2016; Lee *et al*., 2019, 2020) . SYT1 in collaboration with SYT3 function in maintaining DAG homeostasis at the PM during abiotic stress (Ruiz-Lopez *et al*., 2021*a*). Recent structural analyses have demonstrated that the binding of SYT1 to PM lipids occurs through a Ca^2+-^dependent lipid-binding site and by a site for PIP in response to stress (Benavente *et al*., 2021). Furthermore, CLB1 and SYT6 are Arabidopsis SMP proteins. CLB1 is also localized at ER-PM CS and interacts with SYT1 (Lee *et al*., 2020) (Ishikawa *et al*., 2020*a*). And SYT6 has been described to localize in both the ER and Golgi apparatus (Cabanillas *et al*., 2018)

Finally, an early search for Syt-like sequences from the nucleotide sequence databases at NCBI in plants was performed in 2007 (Craxton, 2007). This analysis allowed the initial identification of several other plant proteins with a SMP domain. Here, we have used HHpred server (Zimmermann *et al*., 2018) for a deep search of the remote SMP protein orthologs in Arabidopsis and tomato (*Solanum lycopersicum)* proteomes. Then, we have used confocal microscopy to uncover the subcellular localization of the plant SMP proteins that have not been previously studied. We are reporting the first lipid-transfer proteins localized at ER-chloroplast CS. In this work, we found that NTMC2T5 proteins are anchored to the outer envelope membrane (OEM) of the chloroplast, and they also bind the ER network. Furthermore, the overexpression of these proteins causes the clustering of chloroplast around nucleus which have been related to a general response to pathogen perception in *Nicotiana benthamiana* (Ding *et al*., 2019). Additionally, our studies have demonstrated that there are three SMP proteins at ER-TGN CS in plants (SYT6, NTMC2T6 and TEX2). And, we have found out that these proteins are associated with VAMP721 (as SYT5 (Kim *et al*., 2021)) which is a secretory vesicle-localized protein which drive exocytosis in plants, and it is involved in Arabidopsis pathogen immunity. Results shown in this work are the first steps taken to understand the functions and mechanisms of these putative lipid transport proteins located at contact sites in Arabidopsis and tomato cells and which could be important elements for plant stress tolerance, as shown before for SYT1 and SYT3.

## RESULTS AND DISCUSSION

### Identification of SMP containing proteins in Arabidopsis and tomato

In order to search for SMP containing proteins in Arabidopsis and tomato proteomes, we used the SMP amino acid sequence of the *Homo sapiens* E-syt1 protein (D135-V313) as query. We searched for remote orthologs in all Arabidopsis and tomato proteins using the HHpred server (Zimmermann *et al*., 2018), that unlike PSI-BLAST contemplate equivalent amino acids. As a control, a search using the human proteome (4 Jul 2017) was also carried out and returned all six known SMP-containing proteins in *Homo sapiens* (HsE-Syt1 to 3, HsTEX2, HsPDZD8 and HsTMEM24). The search on the Arabidopsis proteome (TAIR10, 20 Jun 2017) identified a total of fifteen proteins (AtSYT1 to 6, AtCLB1, AtNTMC2T5.1 and 5.2, AtNTMC2T6.1 and 6.2, AtTEX2A and 2B, AT3G60950 and AT3G61030) with a strong confidence in containing a SMP domain. The search in the *S. lycopersicum* proteome (28 Jul 2019) identified sixteen proteins: SlSYT1 (*Solyc01g111520*), SlSYT3 (*Solyc09g007860*), SlSYT5 (*Solyc07g007680*), a protein annotated as Tricalbin3 (*Solyc04g008620*) plus twelve uncharacterized proteins encoded by *Solyc01g094030, Solyc03g019770, Solyc03g119440, Solyc05g047710, Solyc06g072110, Solyc07g061950, Solyc08g021940, Solyc10g017590, Solyc12g009700, Solyc12g009790, Solyc12g089330 and Solyc12g011420* sequences.

Next, a multiple sequence alignment with the identified SMP proteins from Arabidopsis and tomato together with the six human SMP protein sequences were used to generate a cladogram (Fig. S1). Based on these results, we named some of the uncharacterized tomato proteins after their Arabidopsis putative orthologs, and we renamed the tomato Tricalbin 3 protein as SlNTMC2T5 (as it grouped with Arabidopsis NTMC2T5 proteins). In the designation of these tomato genes, we also considered their expression profiles in both species as RNA-seq data of these sequences were extracted from TRAVA database (Klepikova *et al*., 2016) (http://travadb.org/) (Fig. 1A and Fig. S2). This analysis revealed that *SYT1* and *CLB1* are the ones with highest expression levels in both species, while some of the newly identified sequences showed very low expression levels, specially *At3g60950* and *At3g61030*.

**Figure 1:**
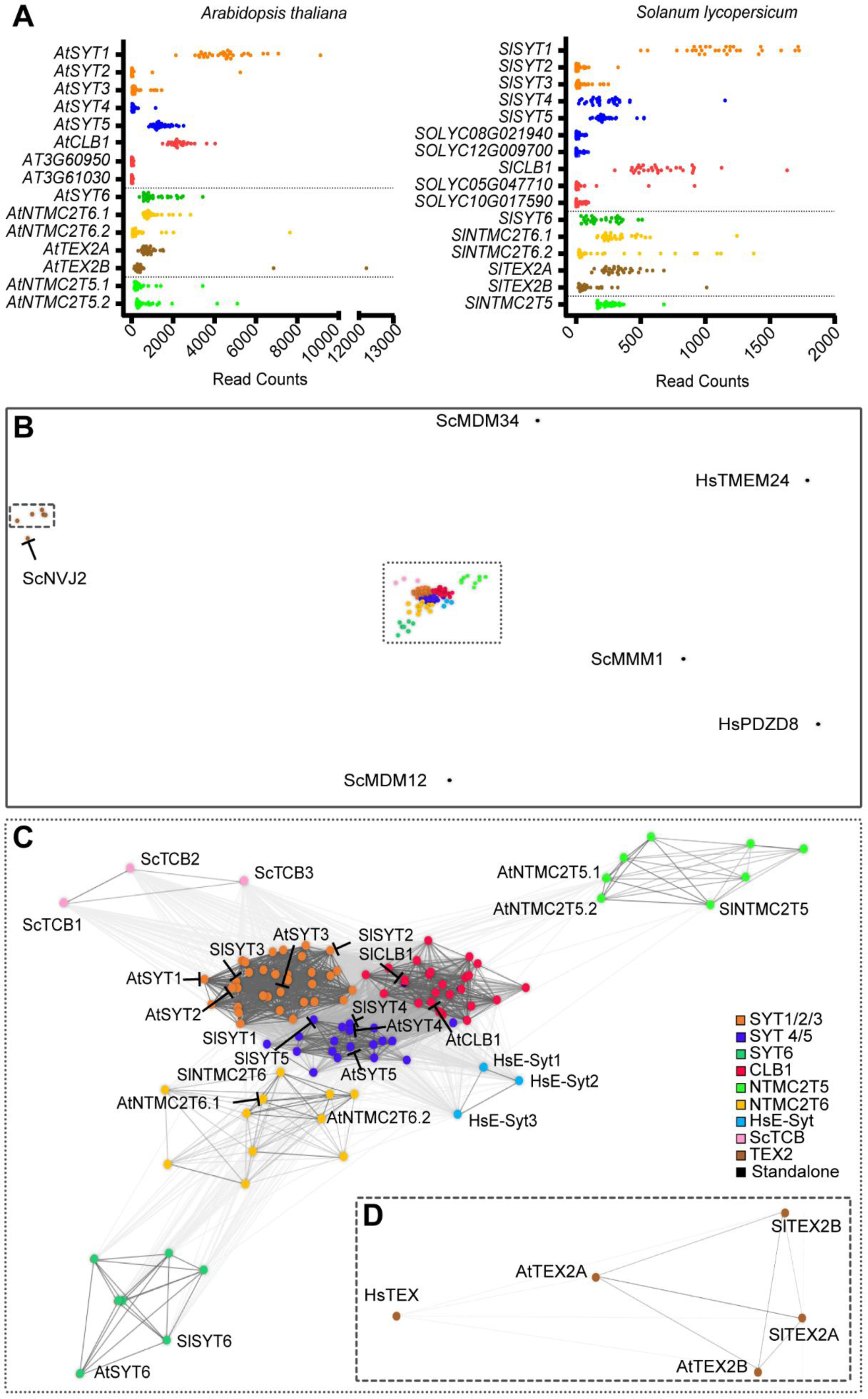
Identification of SMP containing proteins in Arabidopsis and tomato. **(A)** RNA-seq expression data of *A. thaliana* and *S. lycopersicum* proteins was obtained from vegetative tissues at different developmental stages from TRAVA database (http://travadb.org/). Raw norm data was normalized by median-of-ratio method. Each dot represents a value of RPKM. **(B-D)** Clustering map of all SMP containing proteins found in this study plus all proteins listed in (Craxton, 2007) (117 proteins in total) using CLANS (Frickey and Lupas, 2004), based on their all-against-all pairwise sequence similarities. In the map, dots represent sequences and sequences within one group are indicated by the same color. Sequences that could not be assigned to a group are named as standalone proteins. Line coloring reflects the BLAST P-value for their pairwise similarity (brighter color=the lower the P-value). **(C)** Detail of the major cluster of proteins formed by most proteins but HsPDZ8, HsTEM24, ERMES forming proteins from *Saccharomyces* and TEX proteins (details in D).

Then, we clustered all these sequences based on their all-against-all pairwise similarities using CLANS (Frickey and Lupas, 2004). Including all putative SMP protein sequences listed previously (Craxton, 2007) *e.g.* 24 plant SYT-like sequences from *Saccharomyces cerevisiae, Physcomitrella patens* and several plant species (Fig. 1B). In the resulting map the sequences distributed into nine clearly differentiated subfamilies as in the cladogram. Proteins of the ERMES complex from yeast (ScMMM1, ScMDM12, ScMDM34) and human HsTMEM24 and HsPDZ8 proteins did not show any obvious relationship (gray lines) with any of the proteins included in the analysis, and consequently they were tagged as standalone proteins. In the central protein cluster (surrounded by dotted lines), there was a core of highly related proteins that included SYT1-5 and CLB1 plant proteins. Human E-Syt were closer to this central group, and yeast Tcb figured slightly further apart (Fig. 1C). Additionally, three groups of plant exclusive proteins (NTMC2T6, NTMC2T5 and SYT6) showed to be different to the core proteins. Interestingly, we identified a disconnected subfamily of proteins which included TEX2 proteins from human and plants, and yeast ScNvj2 (Fig. 1D). This group of proteins showed some similarity between them, but they were not connected with the rest of analyzed proteins.

### Subcellular localization, protein domains and expression profiles

The prediction of the subcellular localization of Arabidopsis and tomato SMP proteins was performed with DeepLoc 1.0 (Almagro Armenteros *et al*., 2017). Because most MCS take place between the ER and other organelles, ER is expected as the main subcellular location for most SMP-containing proteins. Our results showed that *Arabidopsis* and *Solanum* SYT1 to 5, CLB1, SYT6, NTMC2T6 and three of the four TEX2 proteins were predicted to be localized at the ER (Table S1). Surprisingly, NTMC2T5 proteins were predicted to be localized in the plastid, some putative CLB1-proteins (AT3G60950, AT3G61030, and SOLYC10G017590) were predicted to localize in the mitochondria, SOLYC05G047710 (also within the CLB1 group) was predicted to be a cytosolic protein as it did not show a predicted TM domain, and SlTEX2B was predicted to be a Golgi protein.

SMP proteins usually contain accessory domains to anchor them to the organelle membranes or to regulate their activities. Thus, we also searched for the presence of additional domains in all SMP containing proteins using several bioinformatic prediction tools (*see Materials and method*). We were able to identify transmembrane regions, Pleckstrin homology domains (PH), coiled-coil sequences and FFAT motifs (canonical and FFAT-like) in these proteins. Furthermore, we have analyzed the phosphorylation sites of these proteins (van Wijk *et al*., 2021) and we have studied the predicted three-dimensional (3D) structures of Arabidopsis proteins using several tools, including AlphaFold2 (Jumper *et al*., 2021). These analyses revealed that as it was predicted, all proteins contained a SMP domain. Results from these analyses are summarized in Fig. 2 and in Table S2.

**Figure 2:**
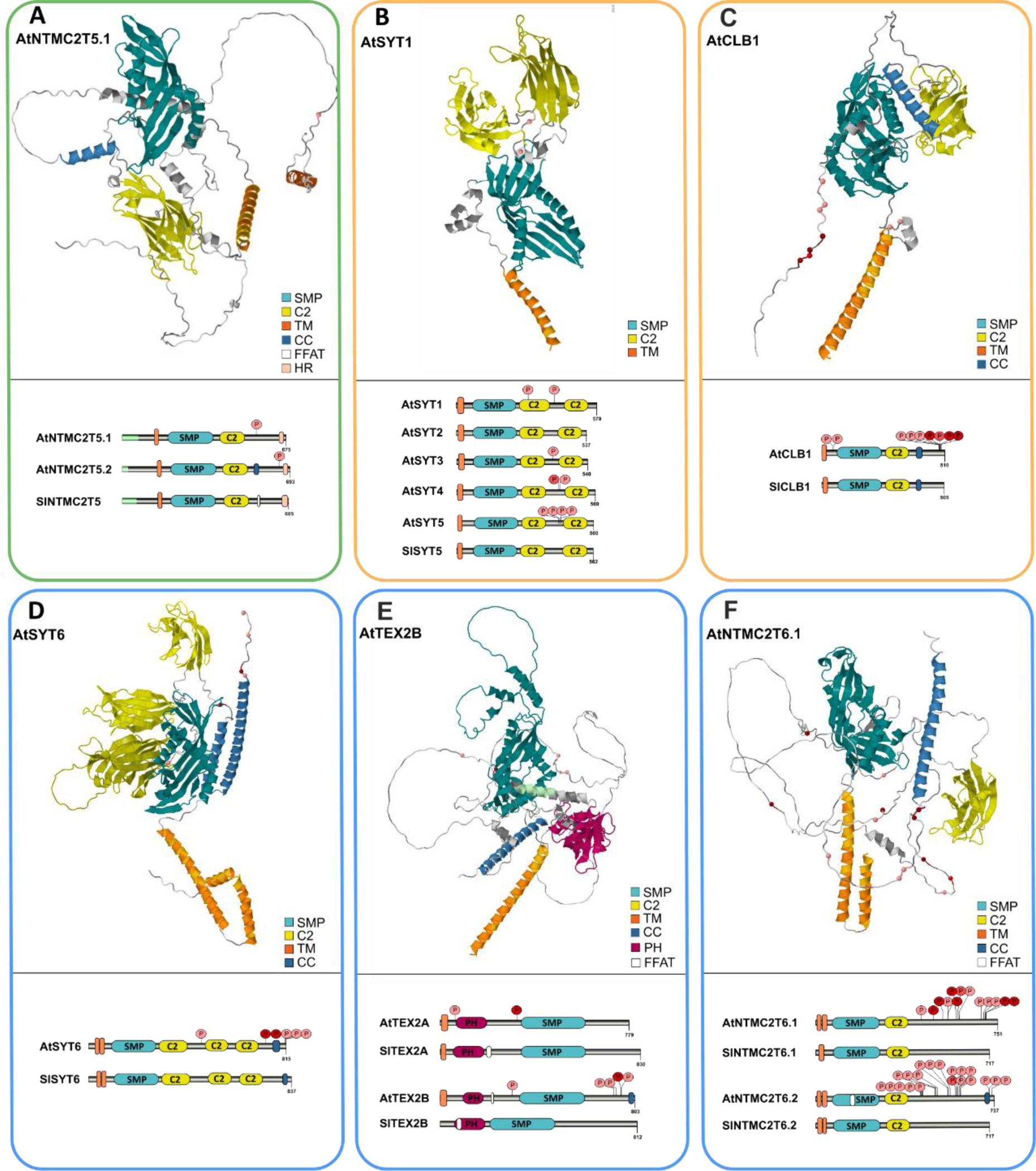
Predicted domains and 3D structures of plants SMP-containing proteins. The structures of *A. thaliana* SMP proteins (one for each subgroup) shown here are derived from AlphaFold (Jumper *et al*., 2021). Domains and motifs were predicted by InterPro, CCTOP and SMART tools. Phosphorylation data was extracted from (Olsen *et al*., 2010)and (Mergner *et al*., 2020). Results are represented using Illustrator for Biological Sequences (IBS) tool (Xie *et al*., 2022) and Jmol. **(A)** NTMC2T5 proteins at ER-Chloroplast CS; **(B and C)** SYT1 to 5 and CLB1 proteins located at ER-PM CS; **(D to F)** SYT6, TEX2 and NTMC2T6 proteins located at ER-TGN CS.

These analyses have revealed the basic structure/domains of all SMP proteins in plants which they already give an insight of the subcellular localization and function of these proteins, however, a much deeper study needs to be carried out. Because SYT and CLB proteins have been previously studied (Kiyosue and Ryan, 1997; Ishikawa *et al*., 2020*b*; Lee *et al*., 2020; Ruiz-Lopez *et al*., 2021*b*), we focused our research on NTMC2T5, SYT6, and NTMC2T6 (all unique to plants) and plant TEX2 proteins (unstudied to date).

### ER-Chloroplast CS: NTMC2T5 proteins

Arabidopsis N-terminal-transmembrane-C2 domain type 5 proteins, NTMC2T5.1 (AT1G50260) and NTMC2T5.2 (AT3G19830), are currently annotated as calcium-dependent lipid-binding (CaLB domain) proteins (source: Araport11). Very limited information is available, apart from the downregulation of these genes in the seedling-lethal deficiency of plastid ATP synthase 1 (*dpa1*) mutant (Dal Bosco et al., 2004). As mentioned above, NTMC2T5 proteins have been predicted to be localized in plastids (Table S1) as they include a chloroplast transit peptide. It was also predicted that they contain a TM, a SMP and a single C2 domain. Additionally, NTMC2T5 proteins are characterized by having a hydrophobic region (HR) at the C-terminal, which is not present in the rest of SMP proteins, and which is predicted to form an alpha-helix of nineteen amino acids in Arabidopsis and twenty-three amino acids in tomato proteins (Fig. 2A).

Thus, we have assessed their subcellular localization, by the transient expression of AtNTMC2T5.1 (as a protein representative of this family) fused to a fluorescent protein in *Nicotiana benthamiana* leaves and by tracking them using confocal microscopy. Ectopic expression of AtNTMC2T5.1, fused to eGFP (green fluorescent protein) at the C-terminus driven by the Ubiquitin10 promoter (UBQ10) from Arabidopsis, showed that this protein is localized around the chloroplasts, identified by the autofluorescence of their chlorophyll (Fig. 3A). Interestingly, chloroplasts with AtNTMC2T5.1-GFP signal tended to clump together. Additionally, we observed a mesh/reticular pattern, suggesting that this protein also localized at the ER. Therefore, we expressed the N-terminus of AtNTMC2T5.1, comprising the predicted signal peptide and its TM domain sequence (M1-G160) followed by eGFP (UBQ10:AtNTMC2T5.1^TM^-GFP). Interestingly, AtNTMC2T5.1^TM^-GFP distributed exclusively throughout the chloroplast membrane (Fig. 3B) supporting that the N-terminal domain of AtNTMC2T5.1 is anchored to chloroplasts.

**Figure 3:**
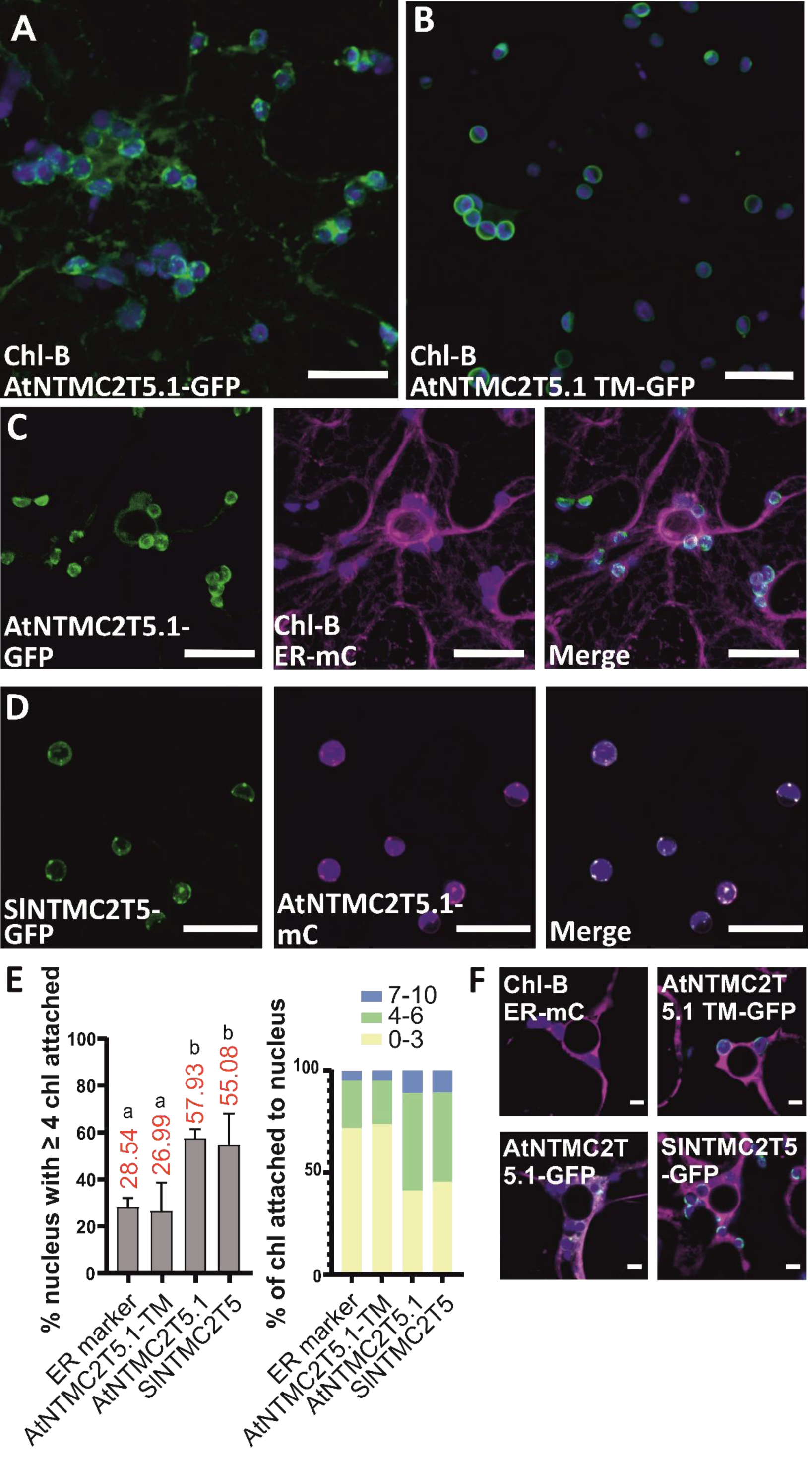
Subcellular localization of NTMC2T5 in *Nicotiana benthamiana*. Transient expression of C-terminus GFP fusion of full-length or truncated versions of AtNTMC2T5.1 protein were analyzed by confocal microscopy (Z-projects shown). **(A)** AtNTMC2T5.1 is surrounding chloroplast (chlorophyll autofluorescence) non-uniformly. AtNTMC2T5.1 also show a mesh/reticular pattern outside chloroplast. **(B)** AtNTMC2T5.1 is anchored to the chloroplast outer membrane, as seen by the expression of its transmembrane domain (M1-G160) fused to GFP. **(C)** Co-expression of full-length AtNTMC2T5.1 with an ER-maker produces clustering of chloroplast surrounded by ER. **(D)** SlNTMC2T5 completely colocalizes with AtNTMC2T5.1. **(E)** Overexpression of full-length AtNTMC2T5.1 and SlNTMC2T5 proteins induces clustering of chloroplast around the nucleus, in contrast to colocalization with the ER marker and AtNTMC2T5.1^TM^-GFP. **(F)** Confocal images of chloroplast clustering around nucleus when expressing the ER-marker on its own, or when coexpressing AtNTMC2T5.1^TM^-GFP or AtNTMC2T5.1-GFP or SlNTMC2T5-GFP constructs together with the ER marker. Scale bar A to D: 20 µm; F: 5 µm.

Next, we co-expressed AtNTMC2T5.1-GFP together with the ER-mCherry marker (CD3-959, signal peptide of AtWAK2 and the ER retention signal HDEL at its C-terminus (Nelson *et al*., 2007)). Confocal images showed clustered chloroplasts surrounded by the ER, indicating that NTMC2T5.1 also contact with the ER (Fig. 3C). Finally, only one copy of this gene was detected in the tomato genome (*SlNTMC2T5*). When *SlNTMC2T5-GFP* was co-expressed with AtNTMC2T5.1-mCherry, they co-localized extensively in the chloroplast OEM, but not homogeneously. Rather, these proteins concentrated at specific regions of OEM. Overall, these results support that AtNTMC2T5.1 is a chloroplast protein which is anchored to the OEM by its TM domain while the cytosolic part is responsible of ER binding, thereby establishing ER-Chloroplasts CS. These data indicated that as expected SlNTMC2T5 and AtNTMC2T5.1 has a similar subcellular localization (Fig. 3D).

Next, we aimed to quantify the chloroplast clustering caused by AtNTMC2T5.1-GFP overexpression (Fig. 3A). For this, we have co-expressed the ER-marker-mCherry with AtNTMC2T5.1^TM^–GFP, AtNTMCT5.1-GFP or SlNTMC2T5-GFP individually. Then, we have quantified the number of chloroplasts around the nucleus in cells of each experiment. The expression of ER marker-mCherry by itself or together with AtNTMC2T5.1^TM^-GFP showed that less than 30% of analyzed nucleus had more than four chloroplasts attached (Fig. 3E and 3F). Interestingly, the expression of AtNTMC2T5.1 and SlNTMC2T5, caused extensive clustering of chloroplasts, as more than 55% of nucleus showed more than four chloroplasts attached. Previous studies have shown that chloroplasts actively cluster around the nucleus following the activation of immune responses (Ding *et al*., 2019), (Mullineaux *et al*., 2020). This clustering has been proved to occur upon perception of different types of invasion patterns or defense-related molecules. Although, our results suggest that NTMC2T5 proteins could be partly responsible of the attachment of chloroplast to the ER surrounding the nucleus, further experiments are needed to confirm this hypothesis and to elucidate the role of these proteins in these processes.

Interestingly, these NTMC2T5 proteins contain a SMP domain which is expected to mediate lipid exchange at MCS. Lipid transport from ER to plastids is understood to occur by non-vesicular transfer at ER-Chloroplasts CS (Wang and Benning, 2012; Hurlock *et al*., 2014; Block and Jouhet, 2015; LaBrant *et al*., 2018; Michaud and Jouhet, 2019). However, it is still an open discussion which lipid(s) are translocated from the ER to plastids or vice versa. To date, PC, lyso-PC, PA and/or DAG have been proposed to be the shuttled lipids from the ER to the chloroplast OEM (LaBrant *et al*., 2018; Karki *et al*., 2019). Thus, as NTMC2T5 proteins contain a SMP domain and characterized SMP domains have shown to transport glycerolipids (AhYoung *et al*., 2015; Reinisch and de Camilli, 2016; Saheki *et al*., 2016*b*; Ruiz-Lopez *et al*., 2021*c*), NTMC2T5 proteins could be responsible of transferring some of the above mentioned glycerolipids involved in the synthesis of chloroplastic lipids. Although, it is likely that more than one mechanism involved in the transport of lipids co-exist in plant cells to ensure the biogenesis of plastid membranes in different conditions, and our data hints NTMC2T5 proteins could be some of them.

### ER-TGN CS: SYT6 proteins

Proteins within SYT1 and SYT5 groups are the most studied proteins, so far. They are characterized by a well-conserved structure with a single TM domain (anchoring the protein to the ER) followed by a SMP and two C2 domains that allow these proteins to contact the PM, Fig. 2B (localizing at ER-PM CS). SYT6 has occasionally been included in the SYT family because SYT6 proteins showed a similar structure to SYT1-5 proteins (Fig. 2D). However, our analyses have found out that they form a separate subfamily of proteins, characterized by having a TM domain likely forming a hairpin (as human E-Syt), followed by a SMP and three C2 domains (as some human E-Syt). Additionally, it is predicted to have a coiled-coil domain formed by two alpha-helix at the C-terminal. Significantly, the Arabidopsis phosphorylation analysis showed one phosphorylation site just before the beginning of the predicted coiled-coil domain and two phosphorylation sites just at the end (S727, S805 and S806) followed by other two phosphorylations at the last end (S812 and S815).

Previous studies using localization of organelle proteins by isotope tagging, LOPIT (Nikolovski *et al*., 2012) showed that AtSYT6 (in addition to AtSYT1, AtSYT5, AtCLB1 and AtTEX2A) was one of the 1,385 proteins targeting Golgi/TGN, but their further classification analysis assigned them into the ER. Later studies showed that the expression of AtSYT6-GFP revealed the presence of some punctate structures which colocalized with a Golgi marker and concluded that AtSYT6 is located both in the ER and the Golgi apparatus (Cabanillas *et al*., 2018). Therefore, we analyzed the subcellular localization of AtSYT6 (AT3G18370) by the transient expression in *N. benthamiana* and confocal microscopy, as before. We have ectopically co-expressed the TM cDNA sequence of AtSYT6 fused to GFP (M1-R83, named AtSYT6^TM^-GFP) driven by UBQ10 promoter with the ER marker-mCherry. Localization of these two protein fragments was highly coincident (Fig. 4A), supporting that AtSYT6 is anchored to the ER by its TM domain, as in SYT1-SYT5 proteins. Then, we have studied the subcellular localization of full AtSYT6 sequence (UBQ10: AtSYT6-GFP, Fig. S3A) and by co-expressing it (Fig. 4B) with a Golgi marker (Man49-mCherry(Nelson *et al*., 2007)). In both cases, AtSYT6-GFP showed the expected reticulated pattern (as it is an ER protein) but some punctuate was also observed. The Man49-mCherry marker labeled the Golgi stacks that mostly appear as round discs, but it also showed weak ER labeling, resulting from the recycling of Golgi resident proteins through the ER (Nelson *et al*., 2007) (Fig. 4B). In our experiments, the dotted AtSYT6-GFP signal did not sufficiently co-localized with the Golgi marker (empty arrows in Fig. 4B), although we noticed some colocalization in some bigger formations of unknown nature. However, as previous proteome reports indicated that AtSYT6 could be localized at Golgi/TGN we hypothesized that AtSYT6 could be predominantly contacting the TGN membranes. Thus, we tested whether AtSYT6-mCherry showed some colocalization with a TGN marker (VAMP721) fused to GFP. VAMP721 is a R-SNARE protein which mainly localizes to the TGN/EE (early endosomes) and the PM and serves as a central hub in post-Golgi trafficking, mediating secretory trafficking in plants (Uemura *et al*., 2019; Shimizu *et al*., 2021). In these experiments (Fig. 4C), we noticed that AtSYT6-mCherry signal was coincident in definite spots of the many small fluorescent VAMP721-GFP vesicles (filled arrows), suggesting the attachment of SYT6 to these vesicles. Lastly, we have studied the subcellular localization of the protein we have identified as the SYT6 ortholog in tomato plants (SlSYT6). Our experiments confirmed that the newly identified SlSYT6 has the same subcellular localization as AtSYT6 (Fig. 4D), as expected both proteins showed a perfect co-localization. So far, our experiments suggest that SYT6 are proteins whose TM is anchored to the ER and the rest of the protein remains in the cytosol or primarily interacting with the TGN forming MCS between these two membranes.

**Figure 4:**
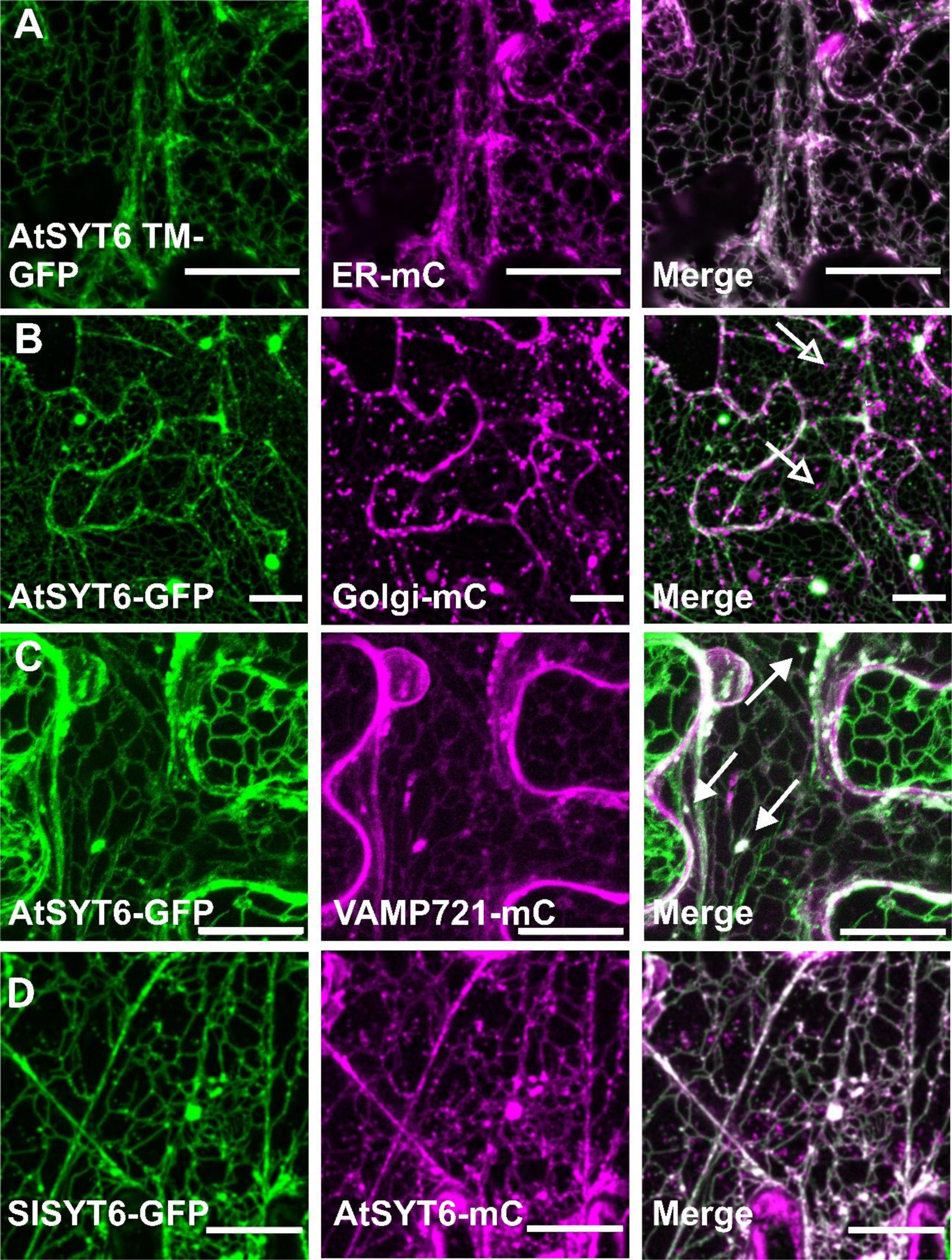
Subcellular localization of SYT6 *in planta*. Transient expression in *N. benthamiana* leaves of C-terminus GFP fusions of full or truncated versions of SYT6 protein were analyzed by confocal microscopy (Z-projects shown). **(A)** AtSYT6 is a protein anchored to the ER, as seen by co-expression of the transmembrane domain (M1-R83) fused to GFP with an ER marker. **(B)** Full-length AtSYT6 shows an ER pattern alongside some punctuate that mostly do no colocalize with the Golgi marker (Man49) but in some bigger formations of unknown nature. Empty arrows point to mismatches between the Golgi marker and AtSYT6 localization. **(C)** Full-length AtSYT6 shows a high degree of colocalization with the TGN/Early endosomes marker VAMP721 (full arrows), suggesting that AtSYT6 is anchored in the ER membrane but interacting with TGN. **(D)** SlSYT6 completely colocalizes with AtSYT6. Scale bar: 20 µm.

### ER-TGN CS: TEX2 proteins

On a step forward, we decided to study the subcellular localization of AtTEX2 proteins in plants. AtTEX2A (AT1G73200) and AtTEX2B (AT1G17820) are putative orthologs of Nvj2p and HsTEX2 and these two proteins promote the transfer of ceramides from the ER to the Golgi complex (Liu *et al*., 2017). As mentioned above, LOPIT analysis also detected AtTEX2A as a Golgi/TGN associated protein. Our analysis on TEX2 sequences revealed the existence of a TM in both AtTEX2 proteins and in SlTEX2A which is consistent with their predicted localization to ER tubules (Table S1). Additionally, it was predicted that these proteins do not contain any C2 domains but a PH domain (which would bind phosphatidylinositol lipids) ahead of their SMP domains and the predicted SMP domains were substantially larger (262 and 271 amino acids in SlTEX2A and SlTEX2B, respectively) than the rest of analyzed SMP domains (usually between 182 and 194 amino acids) (Fig. 2E). Most of analyzed TEX2 sequences contained a FFAT motif next to these PH domains. These FFAT motifs are frequently present in lipid transfer proteins such as oxysterol-binding protein (OSBP), ceramide transfer protein (CERT), or Nir2/3 and many related homologs in human and yeast (Loewen *et al*., 2003; Kaiser *et al*., 2005; Kawano *et al*., 2006). Finally, TEX2 proteins were predicted to contain a coiled-coil domain, suggesting that these proteins could be involved in some transport vesicle tethering (Truebestein and Leonard, 2016).

To further investigate the subcellular localization of AtTEX2 proteins, we expressed the predicted N-terminal TM domain (M1-D50) of AtTEX2B, together with the ER marker (Fig. 5A), as described above. Images showed that AtTEX2B^TM^ is indeed localized at the ER, as SYT6 proteins. In a next step, we have expressed the full protein AtTEX2B fused to eGFP (Fig. S3B) and we co-expressed AtTEX2B-GFP with the Golgi marker (Man49-mCherry), as before. Confocal microscopy images showed that some AtTEX2B-GFP proteins accumulated in certain vesicles/formations, and as opposed to SYT6, we detected some co-localization in a substantial portion of the Golgi marker dotted signals (filled arrows, Fig. 5B). Additionally, when we co-expressed AtTEX2B-GFP with VAMP721-mCherry, AtTEX2B co-localized in most VAMP721 vesicles (Fig. 5C), indicating that AtTEX2B is also anchoring the TGN. Finally, to verify if AtSYT6 and AtTEX2B are colocalizing to some extent, we over-expressed them in *N. benthamiana* leaves (Fig. 5D). The images showed that indeed these two proteins are at the ER where they colocalized, and they mostly (but not completely) overlapped in the Golgi/TGN vesicles (Fig. 5D), agreeing with our previous results which suggested TEX2B seems to be contacting more organelles than TGN (like the Golgi apparatus). Finally, we have co-expressed AtTEX2B-mCherry with SlTEX2A-GFP and these two proteins showed the same localization pattern indicating that the tomato TEX2A has the same subcellular localization as AtTEX2B (Fig. 5E). Overall, these experiments suggest that, even though SYT6 and TEX2 proteins do not have the identical subcellular localization, they both coincided at ER-TGN CS.

**Figure 5:**
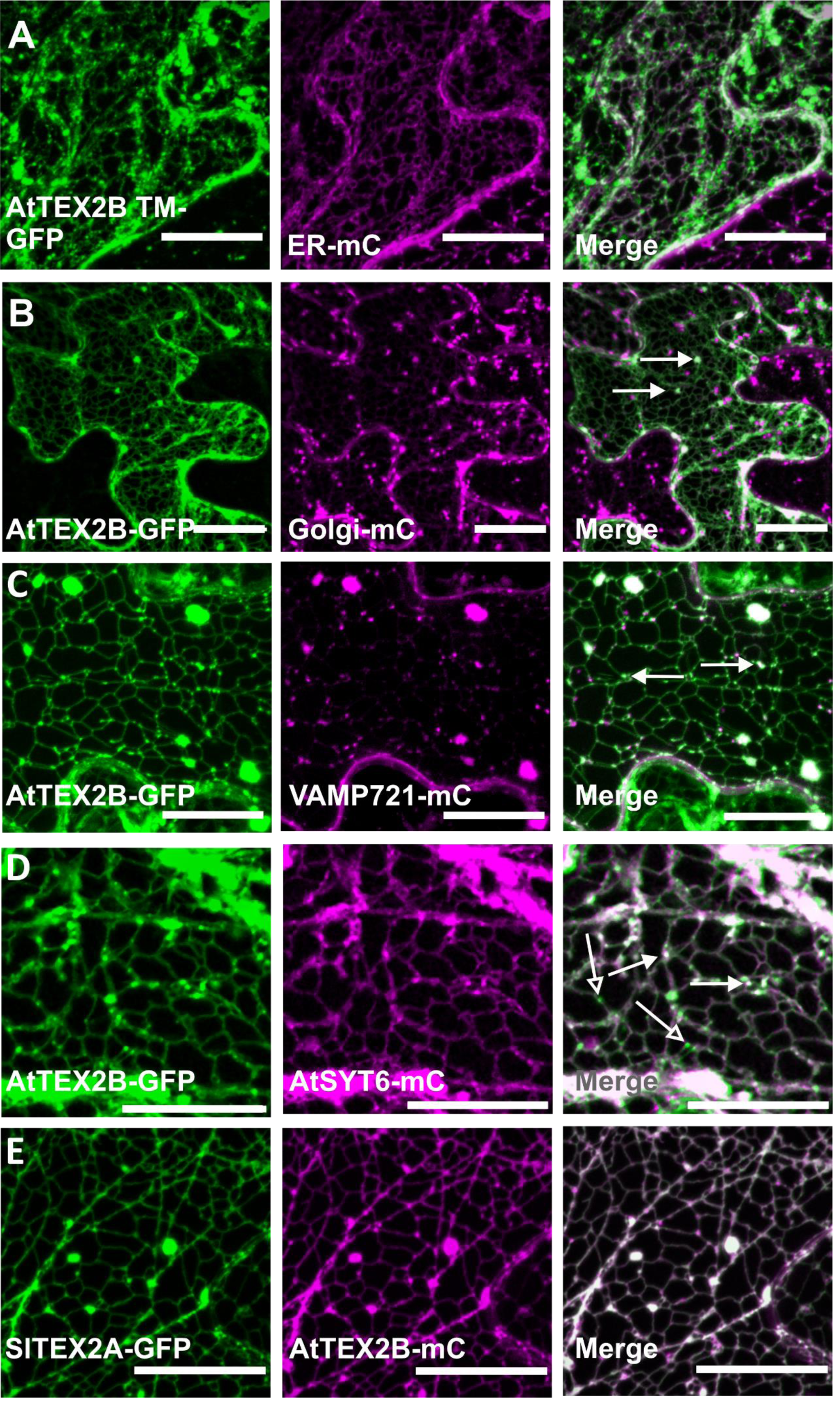
Subcellular localization of TEX2 proteins in *N. benthamiana* leaf epidermal cells. Transient expression of full-length or truncated versions of C-terminally GFP tagged TEX2 proteins were analyzed by confocal microscopy (Z-project shown). **(A)** AtTEX2B is anchored to the ER. Coexpression of transmembrane domain (M1-D50) fused to GFP with ER marker shows colocalization of both signals. **(B)** Full-length AtTEX2B shows a reticular pattern and accumulates at certain vesicles/formations that colocalizes with a portion of Golgi marker Man49 (full arrows). **(C)** Coexpression of full-length AtTEX2B-GFP with the TGN/Early endosomes marker VAMP721 shows almost perfect colocalization indicating that AtTEX2B is interacting with the TGN. **(D)** AtTEX2B-GFP colocalizes with AtSYT6, indicating that both proteins are localized at ER-TGN CS. **(E)** SlTEX2A perfectly colocalizes with AtTEX2B. Scale bar: 20 µm.

### ER-TGN CS: NTMC2T6 proteins

NTMC2T6.1 (AT1G53590) and NTMC2T6.2 (AT3G14590) are plant exclusive SMP proteins, and they have not been studied so far. NTMC2T6.1 has been identified in highly purified vacuoles from mature leaves by 1-D SDS-PAGE LC MS/MS, however it was not present when these fractions were analyzed by 2-D LC MS/MS (Carter *et al*., 2004). It has also been identified as a plasma membrane protein with increased phosphorylation after a flg22 elicitor treatment in Arabidopsis cell cultures (Benschop *et al*., 2007). On the other hand, NTMC2T6.2 has been suggested as a putative plant-specific component of the Arabidopsis ERAD machinery (Lin *et al*., 2019). Therefore, a deeper study on their subcellular location is still pending.

Our *in-silico* analyses have shown that NTMC2T6 proteins share some common structures with SYT6. They are predicted to have two short TM domains in the N-terminal region (except for SlNTMC2T6.1) as shown in Fig. 2F, but their SMP domains are followed by a single C2 domain. These proteins were predicted to have a disorder domain (Fig. 2F) followed by a one helix coiled-coil domain in AtNTMC2T6.2.

Next, we transiently co-expressed the TM of AtNTMC2T6.1 (M1-R73, AtNTMC2T6.1^TM-^GFP) with the ER marker in *N. benthamiana*, as described before. Images showed AtNTMC2T6.1^TM^-GFP as a reticulated pattern which it perfectly co-localized with the ER marker (Fig. 6A), indicating that these are ER anchored proteins. Next, we co-expressed AtNTMC2T6.1 with the Golgi marker. As expected, this protein is located at the ER (Fig. 6B) but, as in SYT6 proteins, the GFP signal is not coincident with the dotted signal of the Golgi marker (empty arrows). We further tested whether AtNTMC2T6.1 is also present in ER-TGN CS as in the case of the SYT6 and TEX2B proteins by co-expressing it with AtVAMP721, AtSYT6 and AtTEX2B. The analysis showed that AtNTMC2T6.1 is highly colocalizing with VAMP721 (Fig. 6C), AtSYT6 (Fig. 6D) and TEX2 (Fig. 6E), although AtSYT6 and TEX2 only share partial colocalization in some regions (domains) of the TGN (filled arrows point similarity in localization, and empty arrows show the differential localization).

**Figure 6:**
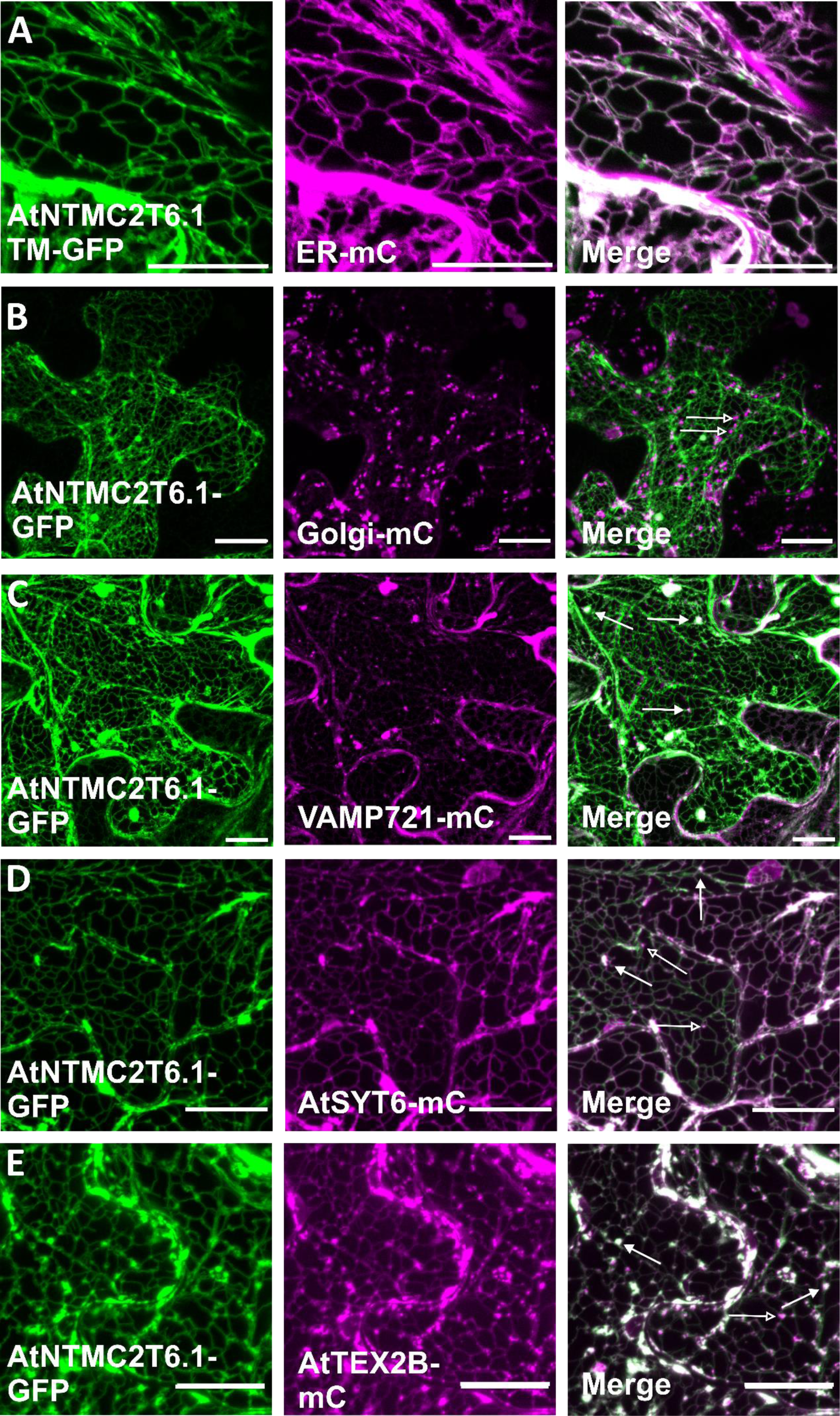
NTMC2T6 subcellular localization. Z-projections of confocal microscopy images of transiently expressing NTMC2T6 in the leaf epidermis of *N. benthamiana* are shown. **(A)** AtNTMC2T6.1 is anchored to the ER, as seen by coexpression of its transmembrane domain (M1-R73) fused to GFP with an ER marker. **(B)** Full-length AtNTMC2T6.1 shows an ER pattern along with some punctuate structures. Abundant mismatches between the Golgi marker and AtNTMC2T6.1 (empty arrows) are also observed. **(C)** Full-length AtNTMC2T6.1 shows a high level of colocalization with the TGN (Early endosomes marker VAMP721) (full arrows), suggesting that AtNTMC2T6.1 is anchored in the ER membrane but interacts with the TGN at ER-TGN CS. **(D-E)** AtNTMC2T6.1 colocalizes with AtSYT6 and AtTEX2B in some areas (filled arrows) but not completely (empty arrows). Scale bar: 20 µm.

Overall, these analyses have shown that SYT6, TEX2 and NTMC2T6 proteins are anchored to the endoplasmic reticulum by their N-terminus TM domains. Furthermore, these proteins showed certain colocalization with each other and with the TGN protein VAMP721, supporting that these proteins function as tethers of ER-TGN CS.

### Association of SYT6, TEX2 and NTMC2T6 proteins at ER-TGN CS

Thus, proteins under study are ER proteins that bind (likely by their C2/PH domains) to TGN vesicles. To confirm this hypothesis, we used Brefeldin A (BFA), an inhibitor of the ARF-GEF (guanine nucleotide exchange factors for ARF GTPases), arresting vesicle trafficking along the secretory pathway. In response to BFA, TGN, endosomal, and most Golgi material aggregate in large intracellular bodies near the nucleus (BFA compartments) (Klausner *et al*., 1992; Nebenführ *et al*., 2002; Langhans *et al*., 2011). The R-SNAREs VAMP721 protein was used as a positive control of the treatment as it has previously been shown these proteins accumulate in BFA bodies after a BFA treatment (Zhang *et al*., 2011, 2021). Additionally, we used a chimeric protein named MAPPER as a negative control. MAPPER is a protein that anchors to the ER and due to its phosphoinositide-binding polybasic motif shows constitutive ER-PM tethering in mammalian and plant cells (Chang *et al*., 2013; Lee *et al*., 2020; Ruiz-Lopez *et al*., 2021*c*). Then, we transiently co-expressed SYT6, TEX2B or NTMC2T6.1 (each protein tagged with eGFP) with VAMP721-mCherry. Next, these *N. benthamiana* leaves were re-infiltrated with BFA and analyzed by confocal microscopy. Subcellular localizations of these proteins after BFA or mock treatments are shown in Fig. 7A. In all cases, BFA caused a redistribution of the fluorescent TGN marker (VAMP721-mC) into BFA bodies that were clearly visible around nucleus. Although proteins under study are anchored to the ER, we observed the re-localization of SYT6, TEX2B and NTMC2T6.1 proteins into BFA bodies, as VAMP721. In contrast, MAPPER did not change its ER-PM CS localization. All together these results are indicating that these proteins are indeed contacting the TGN vesicles.

**Figure 7:**
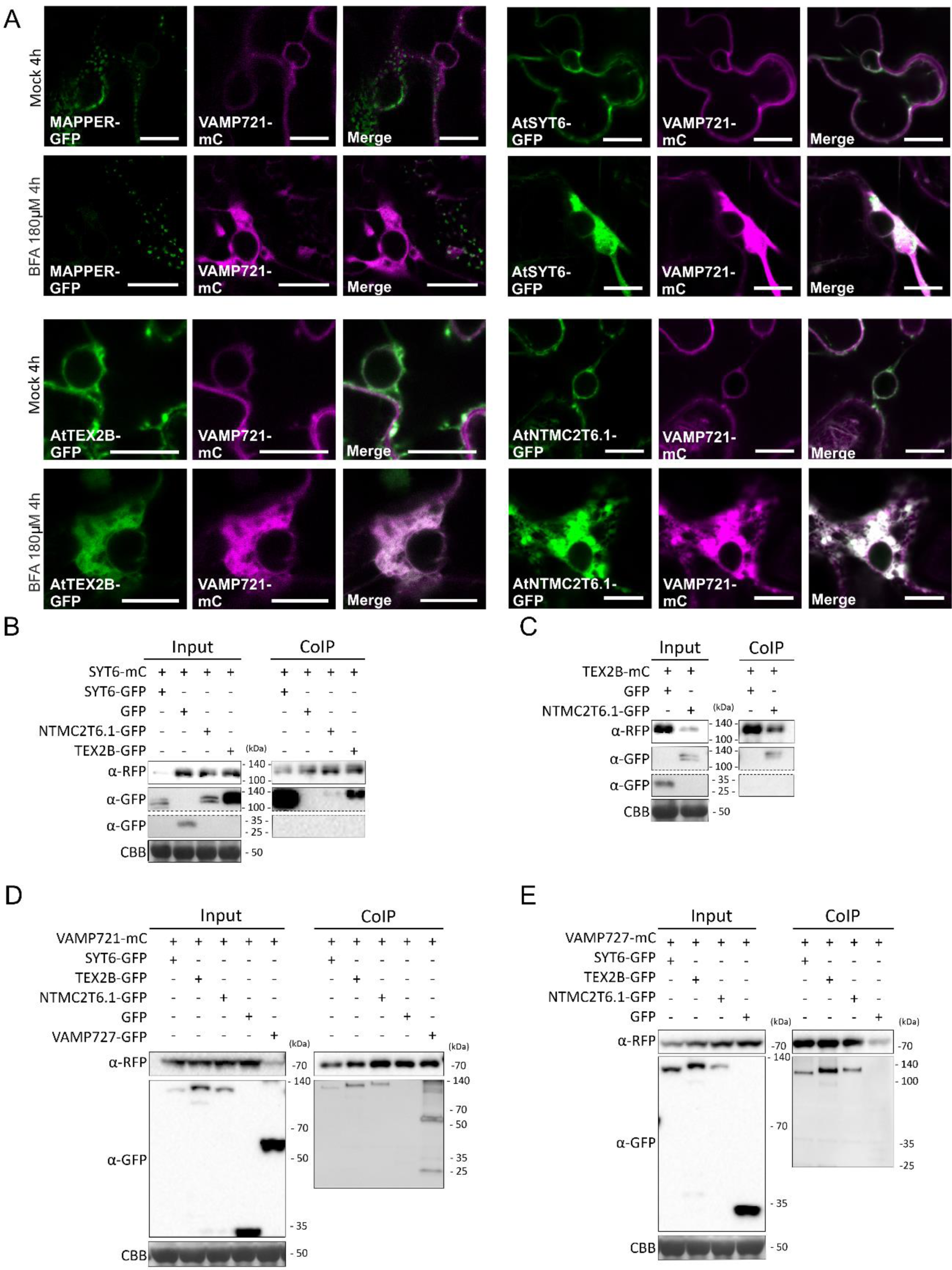
SYT6, NTMC2T6.1 and TEX2B association within ER-TGN CS. **(A)** Relocalization of AtSYT6, AtTEX2B and NTMC2T6.1 into Brefeldin A (BFA) bodies after BFA treatment. Confocal images of *N. benthamiana* epidermal cells transiently expressing AtSYT6, AtTEX2B or AtNTMC2T6.1 fused to GFP with VAMP721-mCherry after BFA treatment (180 µM) or DMSO (Mock). VAMP721-GFP (positive control) shows relocalization into BFA bodies. MAPPER-GFP (negative control) did not relocalize after BFA treatment. **(B)** AtSYT6 coinmuniprecipitates with itself, AtNTMC2T6.1 and AtTEX2B, but not with soluble GFP (negative control). AtSYT6-mCherry was transiently coexpressed with AtSYT6-GFP, AtNTMC2T6.1-GFP, AtTEX2B-GFP and soluble GFP. AtSYT6-mCherry was immunoprecipitated with anti-RFP TRAP beads. Total (input) and Co-Immunoprecipitated (CoIP) proteins were analyzed by western blotting. Equal loading was confirmed by Coomassie blue staining (CBB) of input samples. **(C)** AtTEX2B coimmunoprecipitates with AtNTMC2T6.1. AtTEX2B-mCherry was transiently coexpressed with AtNTMC2T6.1-GFP or soluble GFP (negative control) in *N. benthamiana* and analyzed as described in (B). **(D)** AtSYT6, AtTEX2B, AtNTMC2T6.1 coinmunoprecipitates with VAMP721. VAMP721-mCherry was transiently coexpressed in *N. benthamiana* with AtSYT6-GFP, AtTEX2B-GFP, AtNTMC2T6.1-GFP, soluble GFP (negative control) and VAMP727-GFP (positive control) and analyzed as described in (B). **(E)** AtSYT6, AtTEX2B, AtNTMC2T6.1 coinmunoprecipitates with VAMP727. VAMP727-mCherry was transiently coexpressed in *N. benthamiana* with AtSYT6-GFP, AtTEX2B-GFP, AtNTMC2T6.1-GFP and soluble GFP (negative control) and analysed as described in (B). In all experiments, AtSYT6-mCherry, AtTEX2B-mCherry, VAMP721-mCherry and VAMP727-mCherry were detected with anti-RFP antibody. AtSYT6-GFP, AtTEX2B-GFP, AtNTMC2T6.1-GFP, soluble GFP and VAMP727-GFP were detected with anti-GFP antibody.

The coincident subcellular localization of these three proteins opened the possibility that AtSYT6, AtNTMC2T6.1 and AtTEX2B proteins could be associated at ER-TGN CS. Thus, we performed *in vivo* co-immunoprecipitation (Co-IP) assays after transient co-expression of mCherry full-length AtSYT6 with AtNTMC2T6.1, AtTEX2B or with AtSYT6, tagged with eGFP in *N. benthamiana* leaves. Soluble GFP protein was used as negative control. After immunoprecipitation of AtSYT6-mC, we detected a strong specific association between AtSYT6-GFP and AtSYT6-mC (Fig. 7B), revealing that AtSYT6 is likely forming homodimers. Additionally, AtNTMC2T6.1-GFP and AtTEX2B-GFP also co-immunoprecipitated with AtSYT6-mC while free GFP did not co-IP. Next, the association between AtTEX2B and AtNTMC2T6.1 was also investigated by performing a similar Co-IP experiment (Fig. 7C). Western blots indicated that AtTEX2B-mC also Co-IP AtNTMC2T6.1-GFP (Fig. 7C), as expected. These results are consistent with the point that these three proteins are forming part of / and associating at ER-TGN CS.

In addition, VAMP721 is a TGN protein which interacts with the protein SYP132 (Syntaxin of plants 132) (Fujiwara *et al*., 2014) which in turn is one of the few predicted interactors for the AtSYT6 protein from the Bio-Analytic Resource (BAR) Arabidopsis Interactions viewer database (Geisler-Lee et al., 2007, updated 2018). Furthermore, it has recently been reported that AtSYT5 controls SYP132-VAMP721/722 interaction for Arabidopsis immunity to *Pseudomonas syringae* (Kim *et al*., 2021). For these reasons, we decided to investigate if AtSYT6, AtTEX2B and AtNTMC2T6.1 proteins were associated *in vivo* with VAMP721 by performing Co-IP assays after their transient co-expression in *N. benthamiana* with VAMP727-GFP or GFP, as positive and negative controls, respectively. As previously reported, we detected an association of VAMP721 with VAMP727-GFP (Fig. 7D). Interestingly, we also detected association between VAMP721 and SYT6, NTMC2T6.1 and TEX2B proteins but not with the negative control, indicating that these proteins could be likely forming a complex at ER-TGN CS. Next, we studied the association of these proteins with VAMP727, another R-SNARE at TGN vesicles, that is required for trafficking of storage proteins to the protein storage vacuoles (PSV) (Ebine *et al*., 2008) and SYP41, a Qa-SNARE protein at TGN vesicles. SYT6, TEX2B and NTMC2T6.1 proteins associated with VAMP727 (Fig. 7E), but NTMC2T6.1 proteins did not associate with SYP41 (Fig. S3D). The TGN is an independent membrane compartment on the *trans*-side of Golgi stacks and recent multicolor high-speed and high-resolution spinning-disk confocal microscopy approaches have revealed that TGN proteins exhibit spatially and temporally distinct distribution (Shimizu *et al*., 2021). This could explain why these proteins are all associating with VAMP721 or VAMP727 but NTMC2T6.1 did not associate with SYP41.

All together these results show that SYT6, TEX2B and NTMC2T6.1 are ER proteins, localized at ER-TGN CS forming a complex together with VAMP721 and VAMP727. Furthermore, our confocal and Co-IP analysis suggest that each of them could be contacting specific subdomains of the TGN. The TGN membrane is composed of distinct subdomains which differ in their protein (such as small GTPases, kinases, phosphatases, or lipid transfer proteins) and lipid composition (von Blume and Hausser, 2019). Lipids at the TGN membrane play important roles as they contribute to the recruitment of cargo proteins, to the vesicle fission and to local membrane curvature or tubule formation(von Blume and Hausser, 2019). Thus, these proteins could be functioning as tethers at ER-TGN CS and based on previous studies it is tempting to speculate that AtTEX2 could be transferring ceramide molecules between ER and TGN (as HsTEX2 or Nvj2) and AtSYT6 could be transferring specific glycerolipids (as AtSYT1 or AtSYT3, (Ruiz-Lopez *et al*., 2021*b*)).

Due to the importance of membrane contact sites in cell biology and the limited knowledge, we performed this study to identify proteins with SMP domains in Arabidopsis and tomato. This domain is involved in the transfer of lipids between membranes, and, importantly, all SMP proteins identified to date are located at CS. In this work, we searched for putative sequences encoding for SMP proteins. We have identified NTMC2T5 proteins localized at ER-chloroplast CS. These proteins are extremely interesting, since to the best of our knowledge, proteins involved in lipid crosstalk between chloroplasts and the ER have not yet been identified. Their role in chloroplast clustering around nucleus make them also specially interesting. Our studies showed that SYT6, NTMC2T6 and TEX2 proteins are located at ER-TGN CS, most likely contacting different subdomains of the TGN. Thus, our study provides a starting point for the full characterization on these interesting proteins. Future functional analysis will provide insights into the molecular mechanisms governing lipid traffic between the ER and organelles such as TGN and chloroplasts.

## ABBREVIATIONS

BFA: Brefeldin A
CoIP: Co-Immunoprecipitation
EE: Early Endosomes
eGFP: Enhanced Green Fluorescent Protein
ER: Endoplasmic reticulum
GI-TGN: Independent TGN
IEM: Inner Envelope Membrane
mC: mCherry
(M)CS: (membrane) contact sites
NTMC2(T): N-terminal-TM-C2 (Type) domain proteins
OEM: Outer Envelope Membrane
PH: Pleckstrin homology domain
PM: Plasma membrane
RFP: Red Fluorescent Protein
SMP: synaptotagmin-like mitochondrial-lipid binding SYP132: Syntaxin of plants 132
SYT: synaptotagmin
TEX: Testis-expressed protein
TGN: Trans-Golgi Network UBQ10: Ubiquitin10 promoter

## Supplementary data

**Figure S1:**
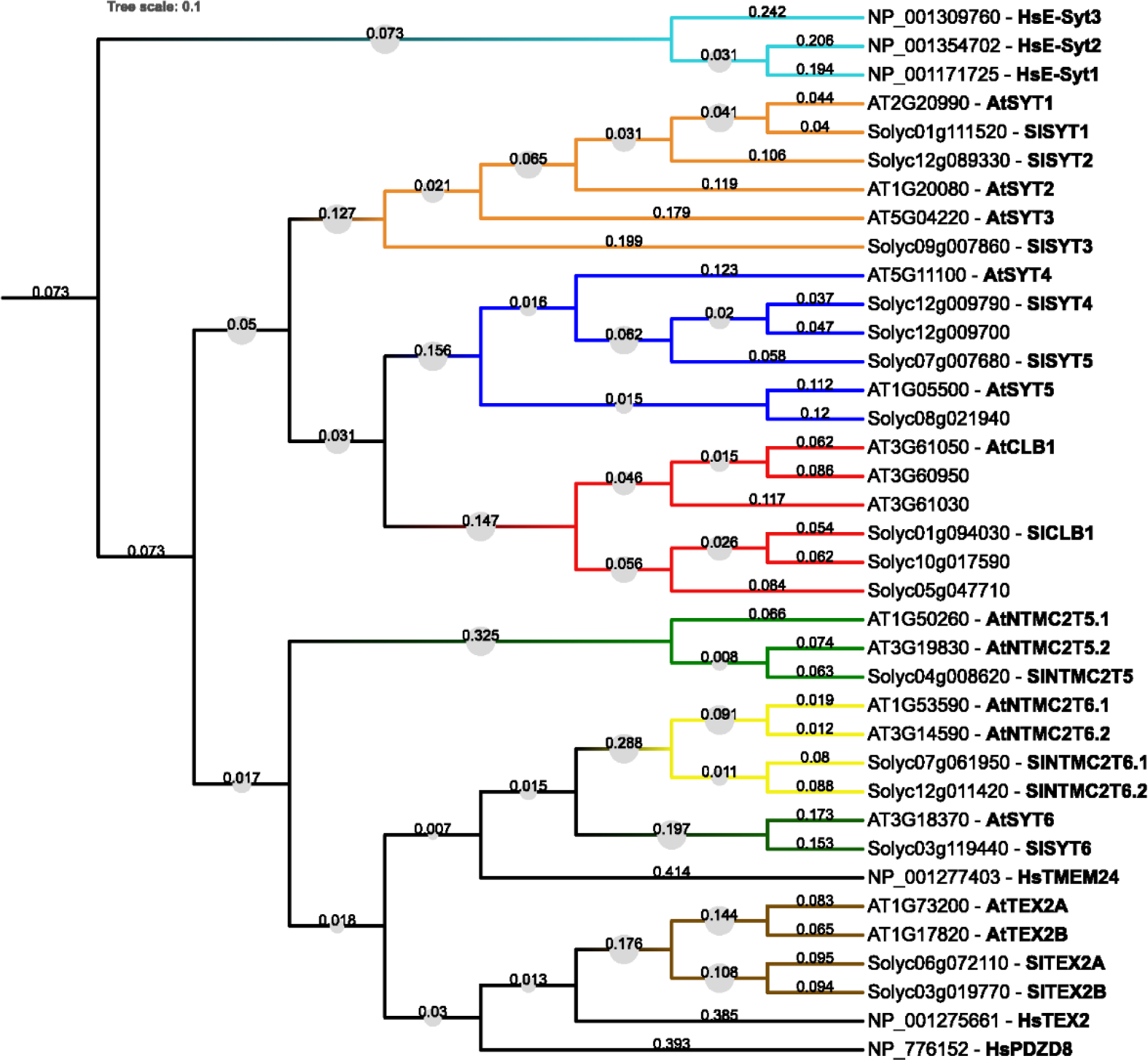
Phylogenetic tree of the *Arabidopsis thaliana*, *Solanum lycopersicum* and *Homo sapiens* SMP-containing proteins. Sequences with high similarity which grouped together with high probability were highlighted in a different color for each cluster. E-Syt proteins from *H. sapiens* were used as outgroup. The tree was constructed using Neighbour-Joining algorithm. Boostrap replications were 500 and p-distance was used as substitution model. The percentage of replicate trees in which the associated taxa clustered together in the bootstrap test is represented with circles over the branches. The number of amino acid differences per site are shown next to the branches (values from 0 to 1).

**Figure S2:**
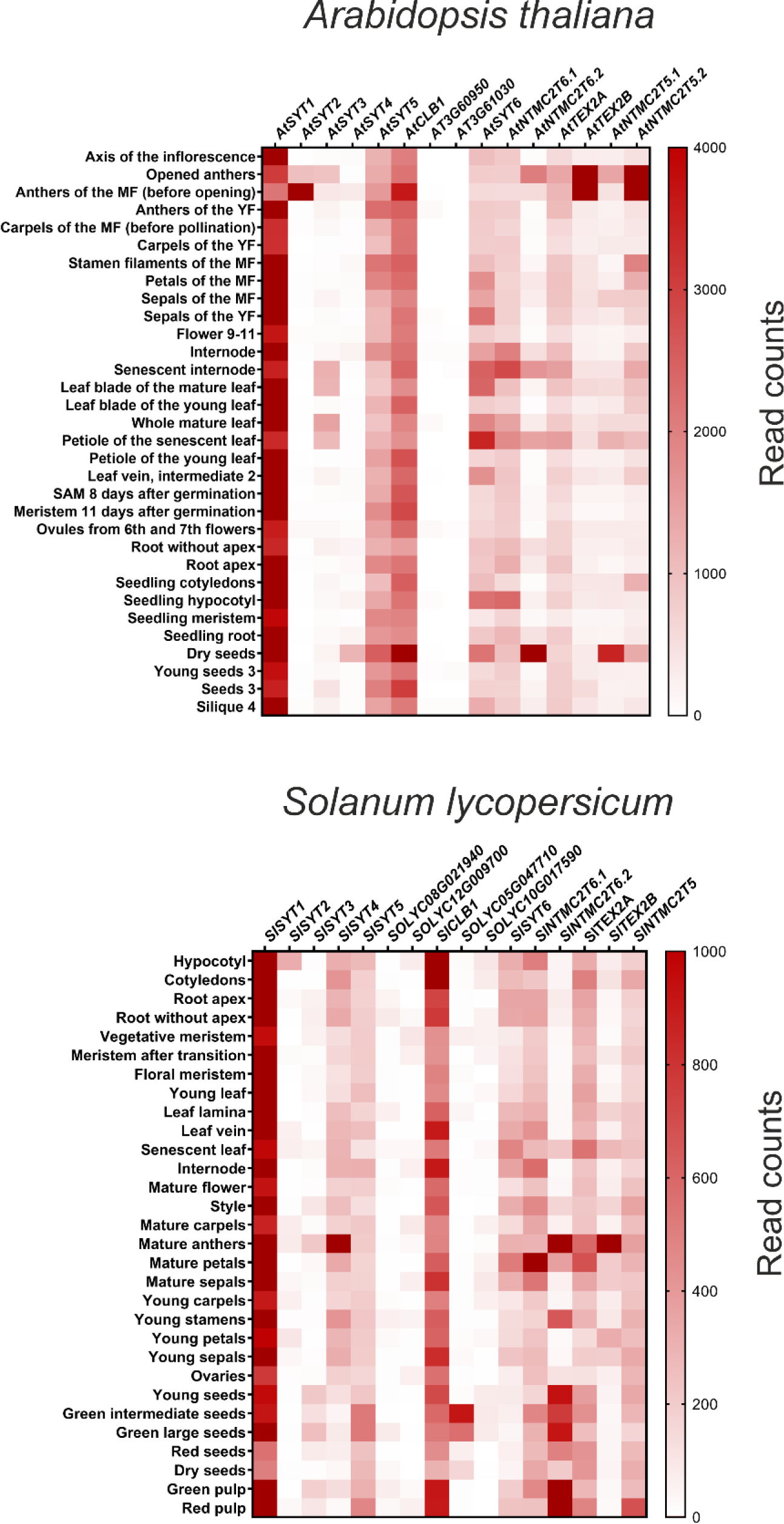
Heat maps of gene expression profiles at different tissues of all SMP proteins in *A. thaliana* and *S. lycopersicum.* RNA-seq data was extracted from TRAVA database (http://travadb.org/). Read-count values were normalized by median-of-ratio method. MF, mature flower; YF, young flower.

**Figure S3:**
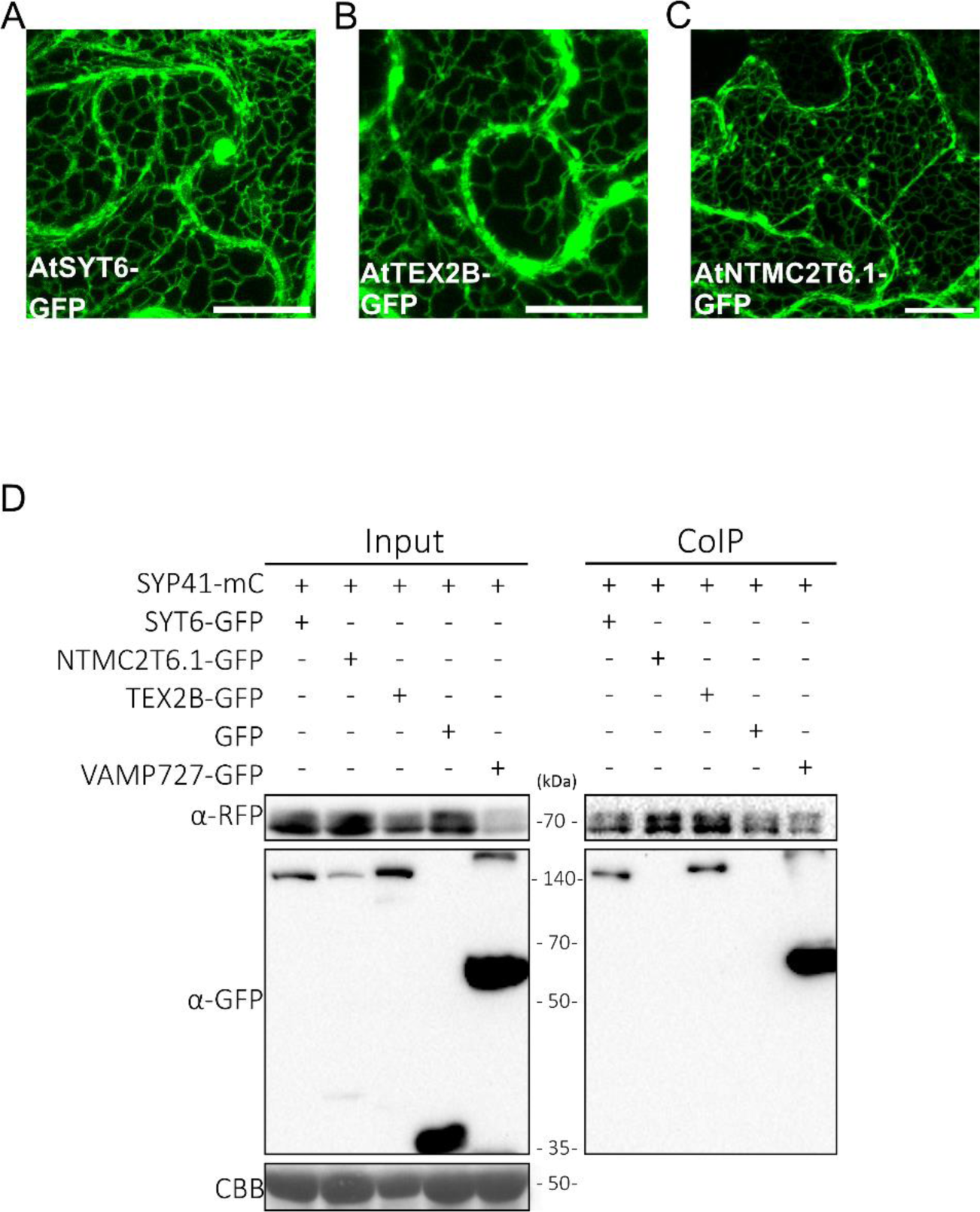
Subcellular localization of AtSYT6, AtTEX2B and AtNTMC2T6.1 in *N. benthamiana* and their association with SYP41. **(A, B, C)** Confocal images of AtSYT6, AtTEX2B and AtNTMC2T6.1 after transient expression in *Nicotiana benthamiana* leaves (Z-projects). All proteins show an ER patter alongside some punctuate structure. Scale bar: 20 µm. **(D)** SYP41 coinmuniprecipitates with AtSYT6 and AtTEX2B, but did not colocalized with AtNTMC2T6.1. SYP41-mCherry was transiently coexpressed with AtSYT6-GFP, AtNTMC2T6.1-GFP, AtTEX2B-GFP, cytosolic GFP (negative control) and VAMP727-GFP (positive control) in *N. benthamiana leaves*. SYP41-mCherry was immunoprecipitated with anti-RFP TRAP beads. Total (input) and co-Immunoprecipitated (CoIP) proteins were analyzed by western blotting. Equal loading was confirmed by Coomassie blue staining (CBB) of input samples. AtSYT6-GFP, AtNTMC2T6.1-GFP, AtTEX2B-GFP, soluble GFP and VAMP727-GFP were detected with GFP antibody and AtSYP41-mCherry was detected with anti-RFP antibody.

Table S1: Predicted subcellular localizations of all SMP proteins in *A. thaliana* and *S. lycopersicum*.

Table S2: Gene name, locus and domains of all SMP proteins found in this study.

Table S3: List of primers used in the present study.

## Acknowledgements

This work was supported by the Ministerio de Ciencia, Innovación y Universidades (PGC2018-098789-B-I00 to N.R-L), by the Ministerio de Ciencia e Innovación (PID2020-120227RJ-I00 to V. S-V and PRE2019-087710 to CH), by the Junta de Andalucía (UMA18-FEDERJA-154 and PAIDI-2020 P20_00222-R to N.R-L, and PAIDI 2020-PY20_00084 to M.A.B). Grant PID2020-114419RB-I00 funded by MCIN/AEI/10.13039/501100011033 and by the “European Union” to MAB.

Additionally, we thank Dr. Eduardo R. Bejaranós lab (IHSM-UMA-CSIC) for providing VAMP721, VAMP727 and SYP41 Agrobacterium strains and Lucía Guirado Manzano and Ana M. López Pérez for their technical support. We thank Alicia Esteban at the microscopy facilities at the IHSM for assistance with confocal microscopy.

## Author contributions

Noemi Ruiz-Lopez conceived the project. All authors contributed to the experimental design. Carolina Huercano and Francisco Percio performed most of the research experiments. All authors analyzed the data and reviewed the manuscript.

## Data availability statement

All data supporting the findings of this study are available within the article and their supplementary materials.

## METHODS AND MATERIALS

### Plasmid constructs/Cloning

The coding DNA sequence (CDS) without the stop codon of *AtNTMC2T5.1* (*AT1G50260*)*, AtSYT6*(*AT3G18370*)*, AtTEX2B* (*AT1G17820*) and *AtNTMC2T6.1* (*AT1G53590*) were PCR-amplified from ArabidopsisCol-0 cDNA, using primer listed in table S3 and cloned into pENTR/ZEO vector by BP cloning kit (Invitrogen). CDS of SlNTMC2T5 (Solyc04g008620), *SlSYT6* (*Solyc03g119440*) and *SlTEX2A (Solyc06g072110*) were PCR-amplified from *S. lycopersicum var Moneymaker* cDNA and cloned as Arabidopsis ones. The transmembrane truncated versions of *AtNTMC2T5.1* (M1 to G160), *AtSYT6* (M1 to R83)*, AtTEX2B* (M1 to D50) and *AtNTMC2T6.1* (M1 to R73) were PCR amplified using the primers detailed in table S3 and built into pENTR/ZEO vectors. All pENTR/ZEO were verified by sequencing. The final Gateway-expression vector was constructed by LR-reaction (Invitrogen). The pENTR vectors, pEN-L4-pUBQ10-R1 and pEN L2-mCherry-R3 or pENT L2-eGFP-R3 were used with the pDEST vector pH7m34GW. Golgi apparatus marker, VAMP721-GFP, VAMP721-mCherry, VAMP727-GFP and SYP41-GFP expression vectors were a gift from Dr. Eduardo R. Bejarano Lab’s. Cis-medial Golgi marker (Man49-mCherry) and ER marker (CD3-959) were retrieved from (Nelson *et al*., 2007).

### Transient expression in *N. benthamiana*

Transient expression in *N. benthamiana* leaves was performed using *Agrobacterium* strain GV3101::pMP90 carrying the appropriate constructs, together with the A*grobacterium* line expressing the silencing suppressor p19. Plants were grown for 21 days at long day conditions, 24 ⁰C, 65% of humidity and 120 µmol photon m^−2^ s^−1^. *Agrobacterium* cultures were grown overnight in Luria-Bertani medium containing rifampicin (50 mg mL^-1^), gentamycin (25 mg mL^-1^) and the construct-specific antibiotic. After growth, bacteria were harvested by centrifugation and pellets were incubated with infiltration solution (10 mM MES pH 5.6, 10 mM MgCl_2_, and 1 mM acetosyringone) for 2 h in dark conditions at room temperature. For single construct experiments, *Agrobacterium* (transformed with the appropriate construct) was infiltrated at OD_600_ of 0.6 and of 0.3 for the p19 strain. For double infiltration experiments, *Agrobacterium* strains were infiltrated at OD_600_ of 0.4 for the constructs and 0.2 for the p19 strain. *N. benthamiana* leaves were infiltrated with *Agrobacterium* into at the abaxial side of the leaf lamina. After infiltration, all plants were kept in the growth chamber, at conditions described above and analyzed at 2 days post infiltration.

### BFA treatment

For BFA (Brefeldin A) treatments, constructs to test were agroinfiltrated in *N. benthamiana* leaves as detailed above. BFA solutions in DMSO were prepared at a final concentration of 180 µM diluted in water. Leaves were infiltrated with BFA solution using a syringe 4 hours before confocal analysis. Mock leaves were infiltrated with 0.36 %(v/v) DMSO diluted in water (same DMSO concentration than in BFA treatment).

### Confocal imaging

For imaging on *N. benthamiana* leaves, 2 days after infiltration leaf-disks were excised from the leaves immediately before visualization under the LMS 800 confocal microscope with the objective C-Apochromat 40x/1.2 W Korr M27. The lower epidermis of the leaf was 3D imaged from the equatorial plane until the cell surface with 1 µm spacing. GaAsp HyD detectors were used to improve the signal detection. For colocalization, sequential line scanning mode was used to separate signals. Cortical plane images are a maximum Z-projection of several planes from the cell surface until a plane where cells are close but still not touching the neighbors (to ease identification of individual cells). Equatorial plane images are single plane images.

### Image analysis

All images were analyzed using FIJI (Schindelin *et al*., 2012).

### Protein identification and dendogram/cladogram representations

Sequences were retrieved from NCBI using their accession number, provided by HHPred (Zimmermann *et al*., 2018) using n° of target sequences 10000, Min. probability in hit list 50% and using HsESyt1 as query. Alignment was carried out in the online version of MAFFT (https://mafft.cbrc.jp/alignment/server, Standard settings). The resulting dendrogram/cladogram was developed in MEGAX (Ver. 10.1.7) with Neighbor-Joining algorithm (Boostrap replications: 500, Substitution model: p-distance). The percentage of replicate trees in which the associated taxa clustered together in the bootstrap test are shown next to the branches (Values from 1 to 100). The number of amino acid differences per site are also shown next to the branches (values from 0 to 1). Sequences with high similarity which grouped together with high probability were highlighted in a color for each cluster.

### CLANS

SMP protein sequences found in this study were clustered together with all putative SMP sequences found in Craxton 2007 (Craxton, 2007) from a wide variety of species (*e.g. Saccharomyces cerevisiae*, *Physcomitrella paten*s and several plant species). Clustering was developed using the CLANS tool (Frickey and Lupas, 2004) from MPI Bioinformatics Toolkit (Scoring Matrix: BLOSUM62, E-value threshold: 1e^-9^) and visualized using the Java executable version of CLANS (standard settings).

### *In silico* analysis

RNA-seq expression data from TRAVA database (http://travadb.org/) was used to determine the gene expression profiles. Different developmental stages of *A. thaliana* and *S. lycopersicum* were analyzed. Normalized raw norm data are represented in a heatmap. Normalization was done by median-of-ratio method. The resulting data was represented in GraphPad Prism (Ver. 8.0.2).

DeepLoc1.0 (Almagro Armenteros *et al*., 2017) predictor server (based in deep learning) was used to predict the subcellular localization of all SMP proteins found in this study.

For protein sequence analyses InterPro (http://www.ebi.ac.uk/interpro/) was used for the prediction of all protein domains, except for transmembrane domains that were predicted using CCTOP tool (Dobson *et al*., 2015). Also, SMART tool (Letunic *et al*., 2021) was used to search for PH domain and the Coiled-coil server (Lupas *et al*., 1991) to search for coiled-coil regions. Proteins were further modelling using Phyre2 (Kelley *et al*., 2015) and AlphaFold2 (Jumper *et al*., 2021).

The presence of FFAT motifs were determined using the ScanProsite tool (https://prosite.expasy.org/scanprosite/). The screening of the FFAT motif was done using the pattern [DE]-X(0,5)-X-[FY]-[FYCILMVWH]-[DEST]-[ACST]-X-[DESTGNQ] (Mikitova and Levine, 2012).

Experimental data of phosphorylation sites of Arabidopsis and human proteins was collected from (Olsen *et al*., 2010) and (Mergner *et al*., 2020)

### Coinmunoprecipitation (CoIP) of proteins

CoIP experiments were carried out as described in (Amorim-Silva *et al*., 2019) with some minor modifications. Agroinfiltrated *N. benthamiana* leaves were grinded to fine powder in liquid nitrogen. Approximated 0.5 g of grinded tissue per sample were used and total proteins were then extracted with extraction buffer [150 mM Tris–HCl, pH 7.5; 150 mM NaCl; 10% glycerol; 10 mM EDTA, pH 8; 1 mM NaF; 1 mM Na_2_MoO_4_ ·2H_2_O; 10 mM DTT; 0.5 mM PMSF; 1% (v/v) P9599 protease inhibitor cocktail (Sigma)], and 0.5% (v/v) of Nonidet P-40 (CAS: 9036-19-5, from USB Amersham life science) was added at 2 mL/g of powder . Samples were mixed by using an end-over-end rocker for 30 min at 4 ⁰C and centrifuged 20 min at 15,000g (4 ⁰C). Supernatants were filtered by gravity through Poly-Prep Chromatography Columns (#731-1550 Bio-Rad) and 100 µL were reserved for immunoblot analysis as input. The remaining supernatants were diluted (1:1 dilution) in extraction buffer without detergent. The final concentration of detergent (Nonidet P-40) was adjusted to 0.25% (v/v) to avoid unspecific binding to the matrix as recommended by the manufacturer. Protein samples were then incubated for 1.5 h at 4 ⁰C with 25 µL RFP-Trap agarose beads (Chromotek) in an end-over-end rocker. Beads were then collected and washed four times with the extraction buffer. Finally, beads were resuspended in 75 µL of 2x concentrated Laemmli including protease inhibitor cocktail. Samples were heated at 37 ⁰C for 30 min to dissociate immunocomplexes from the beads. Total (input), IP, and CoIP proteins were separated in a 10% SDS–PAGE and analyzed by immunoblot as described below.

### Western Blot analyses

Input, immunoprecipitated and CoIP proteins were separated by SDS–PAGE with (10%) polyacrylamide gels. Proteins were electroblotted using Trans-blot Turbo Transfer System (Bio-Rad) onto polyvinylidene difluoride (PVDF) membranes (Immobilon-P, Millipore) following instructions by the manufacturer. PVDF membranes, containing electroblotted proteins, were then incubated with the appropriate primary antibody followed by the appropriate secondary peroxidase-conjugated antibody. The following primary antibodies were used for detection of epitope-tagged proteins: mouse monoclonal anti-GFP clone B-2 (1:1,000, catalog no. sc-9996, Santa Cruz Biotechnology) and anti-RFP antibody (1/2000, Chromotek 6G6). Secondary antibody anti-mouse IgG whole molecule-Peroxidase (1:80,000; catalog no. A9044, Sigma-Aldrich) was used. Proteins and epitope-tagged proteins on immunoblots were detected using the Clarity ECL Western Blotting Substrate or SuperSignal West Femto Maximum Sensitivity Substrate according to the manufacturer’s instructions, and images of different time exposures were acquired using the Chemidoc XRS1System (Bio-Rad). SDS–PAGE and immunoblotted PVDF membranes were stained with Coomassie Brilliant Blue R 250 to visualize equal loading of the different samples.

